# Dynamic Prediction During Perception of Everyday Events

**DOI:** 10.1101/348946

**Authors:** Michelle L. Eisenberg, Jeffrey M. Zacks, Shaney Flores

## Abstract

The ability to predict what is going to happen in the near future is integral for daily functioning. Previous research suggests that predictability varies over time, with increases in prediction error at those moments that people perceive as boundaries between meaningful events. These moments also tend to be points of rapid change in the environment. Eye tracking provides a method for continuous measurement of prediction as participants watch a movie of an actor performing a series of actions. In two studies, we used eye tracking to study the time course of prediction around event boundaries. In both studies, viewers looked at objects that were about to be touched by the actor shortly before the objects were contacted, demonstrating predictive looking. However, this behavior was modulated by event boundaries: looks to to-be-contacted objects near event boundaries were less likely to be early and more likely to be late, compared to looks to objects contacted within events. This result is consistent with theories proposing that event segmentation results from transient increases in prediction error.

**Significance Statement:** The ability to predict what will happen in the near future is integral for adaptive functioning, and although there has been extensive research on predictive processing, the dynamics of prediction at the second by second level during the perception of naturalistic activity has never been explored. The current studies therefore describe results from a novel task, the Predictive Looking at Action Task (PLAT) that can be used to investigate the dynamics of predictive processing. Demonstrating the utility of this task to investigate predictive processing, this task was applied to study the predictions made by Event Segmentation Theory, which suggests that people experience event boundaries at times of change and unpredictability in the environment. The results of these studies are of interest to communities investigating the dynamic comprehension and segmentation of naturalistic events and to communities studying visual perception of naturalistic activity.

## Introduction

The ability to anticipate what is going to happen in the near future is essential for survival. Prey animals must make predictions about the locations of their predators in order to avoid being eaten. Predators must anticipate the location of their prey so as not to starve. Predictive processing has been shown to play a central role in functions ranging from object recognition (e.g., Bar et al., 2006) to action guidance (e.g., Grush, 2004) to deliberative decision-making (e.g., Doya, 2008). Across these domains, humans and other organisms form representations that correspond to what is likely to happen in the near future.

Predictive processing spans a range of spatial and temporal scales. For everyday human activity, predictions on the temporal scale of a fraction of a second to a few tens of seconds, and on the spatial scale of centimeters to tens of meters, are particularly relevant. These are the sorts of predictions that allow people to anticipate whether they can fit their luggage in a car trunk, how a conversation with a friend is going to turn out, or whether a hot dog on the fire will burn.

Previous theories have described the format of representation that might underlie such predictions and the type of computations that might allow people to make predictions in everyday situation. Goodwin and Johnson-Laird (2005), for example, proposed that people typically form a single mental representation of a given situation that corresponds to features of the situations the model represents. Barsalou, Santos, Simmons, and Wilson (2008) extended this theory to language processing and proposed that as people perceive information, they build a multimodal mental representation of all relevant features and the context in which the information was perceived. When this information must be recalled at a later date, people reinstate this information as a *situated simulation* that allows people to make inferences and decisions. The Theory of Event Coding (Hommel, 2009) takes mental models one step further, suggesting that there is no difference between a person’s representation of a perceived event and the representation of their own produced actions. Therefore, the representation can be used to both anticipate future actions and carry them out. A major component of all of these theories is that in order to make effective predictions about what is going to occur in the near future, people build mental models, make predictions based on these models, and update these models as needed.

Clearly, prediction is not a static process, and one feature of everyday activity is that predictability varies over time such that at some times predictions are accurate and prediction error is low, whereas at other times prediction errors can spike suddenly. For example, when cooking a hotdog on a fire, it is easy to predict what is going to happen for the first few minutes: the hotdog will slowly become warmer and warmer as it begins to cook. However, at a certain point, the hotdog will quickly begin to turn brown, and it is difficult to predict exactly when it will go from browning to burning, or even catching on fire.

This variability in predictability over time may be reflected in the mechanisms underlying human action control. Norman and Shallice (1980) proposed that human action consists of periods of relatively automatic action with little need for attentional control, interspersed with short periods of active processing when new tasks need to be initiated or when actions that conflict with habitual behavior need to be enacted. More recently, this *contention scheduling* model was tested computationally by Cooper and Shallice (2000), who showed that this model of action selection can produce complex and naturalistic sequences of actions (see also Botvinick & Plaut, 2004). The mechanism of contention scheduling in action performance suggests a parallel in action perception: People may be able to take advantage of the reliable sequential structure in human action to guide their predictions, but at those moments when contention scheduling is needed by an actor, activity will be less predictable to an observer, leading to transient increases in prediction error.

Observers of activity can certainly learn to take advantage of sequential structure—even when that sequentiality is divorced from goal-directed action. Avrahami and Kareev (1994) showed that people could learn to recognize sequences of action that tend to appear together and represent these predictable sequences of activity as events. People use statistical dependences to segment information streams in multiple domains, including language (e.g., Saffran, Newport, Aslin, Tunick, & Barrueco, 1997) and human action (e.g., Baldwin, Andersson, Saffron, & Meyer, 2008). Recognition of sequences is often an implicit process, with observers learning to make predictions based on learned dependencies without having explicit awareness of the sequences (Swallow & Zacks, 2008).

Contemporary models have proposed accounts of how this variability in predictability is used by the perceptual system to allow for adaptive processing of sequences of human activity. For example, one recent theory, Event Segmentation Theory (EST; Zacks, Speer, Swallow, Braver, & Reynolds, 2007), describes the temporal dynamics of predictive processing in everyday event comprehension. EST proposes that people maintain a working memory representation of the current event that informs perceptual predictions about future activity. These predictions are compared with sensory inputs from the environment to calculate a prediction error—the difference between the prediction and what actually occurs. When prediction error rises transiently, this triggers updating of the working memory representation. In naturalistic activity, prediction error increases tend to happen when features in the environment are changing rapidly. For example, when watching someone prepare for a party, people would experience low levels of prediction error as the actor walks around the table and sets plates in front of each seat, because it is easy to predict that the actor is going to continue setting the table. However, when the actor finishes setting the table, there are many possible actions in which the actor could engage, and prediction error would increase as the actor switches to blowing up balloons. This increase in prediction error would cause the event model to update to better represent the actor’s new goal of blowing up the balloons, causing people to experience a subjective event boundary. As the actor continues to blow up balloons, prediction error would decrease and the cycle would begin again.

EST relates the computational mechanism of prediction error-based updating to the subjective experience of events in a sequential stream of behavior. When an event model is updated, the perceiver experiences that one event has ended and another has begun. The subjective experience of event boundaries can be studied using a unitization task, in which participants are asked to press a button whenever they believe one meaningful activity has ended and another has begun (Newtson, 1973). Using boundaries defined with this technique, Zacks, Kurby, Eisenberg, and Haroutunian (2011) tested the hypothesis that event boundaries correspond to points of high prediction error. They asked participants to watch movies of an actor doing everyday activities (i.e., washing a car, putting up a tent). These movies were paused periodically, and participants made predictions about what would occur five seconds later in the movies. The movies were then restarted, and participants received feedback about the accuracy of their predictions by continuing to watch the movies. Some of the pauses occurred right before a boundary between activities, such that the to-be-predicted activity was part of a new event, and some of the pauses occurred within activities, such that the to-be-predicted activity was part of the current event. The authors found that predictions were less accurate when predictions were made across event boundaries than when predictions were made within events. Further, a functional MRI experiment demonstrated that structures in the midbrain associated with signaling prediction error were more activated when participants attempted to predict across an event boundary. These results provide evidence that prediction failures are associated with the perception of event boundaries. However, this study was limited, first, in that comprehension was stopped repeatedly to administer the prediction task and, second, in that it provided very little information about the temporal dynamics of prediction error.

Other relevant data come from studies using a narrative reading paradigm (e.g., Speer & Zacks, 2005; Speer, Zacks, & Reynolds, 2007; Zacks, Speer, & Reynolds, 2009). In these studies, event boundaries tended to be identified at points when many features of the situation were changing, consistent with the suggestion that in naturalistic activity periods of change tend to produce prediction errors. Participants’ reading times slowed at event boundaries, and when readers were asked to rate the predictability of each clause they rated event boundaries less predictable. Pettijohn and Radvansky (2016) showed that editing the text to make a feature change predictable eliminated slowing in reading time, consistent with the idea that event boundaries are associated with spikes in prediction error.

For the visual comprehension of naturalistic everyday activities, eye tracking provides a promising method for studying the time course of predictability. Eye tracking has been used to study predictive looking behavior in people ranging from infants to adults. For example, Hunnius and Bekkering (2010) studied predictive looking in infants using a paradigm in which the infants watched an actor use an object multiple times while the infants’ eyes were tracked using an eye tracker. On some trials, the actor used the object in a typical fashion (e.g., bringing a hairbrush to the head) and, on other trials, the actor used the object in an atypical fashion (e.g., bringing a hairbrush to the mouth). The authors found that infants predictively looked at the location where the object was typically used, even when the actor brought the object to the atypical location, meaning that the infants were not solely using motion information to make these predictions. Predictive looking has also been studied in adults. Flanagan and Johansson (2003), for example, had participants watch an actor move three blocks from one side of the table to the other while their eyes were tracked using an eye tracker. The authors found that participants started looking at the location where the blocks would be moved before the blocks arrived there, suggesting that participants were predicting the block movements. Similarly, Vig, Dorr, Martinez, and Barth (2010) found that when adults watched brief naturalistic scenes, eye movements to salient stimuli were nearly instantaneous, despite the fact that controlled laboratory studies have found that it takes an average of 200 ms to saccade to and fixate on a newly presented stimulus. The authors suggest that participants made predictive eye movements to the locations where salient information would soon be presented.

Predictive looking has also been studied extensively in the context of sports. For example, Hayhoe, McKinney, Chajka, and Pelz (2012) studied predictive eye movements as participants played squash. They found that participants made anticipatory eye movements ahead of the ball’s position at multiple time points throughout the ball’s flight toward them, rather than simply tracking the ball’s actual location. In addition, Diaz, Cooper, Rothkopf, and Hayhoe (2013) used a virtual racquetball task, and found that participants made predictive eye movements to locations above where the ball would bounce and varied their predictive fixations based on the ball’s bounce speed and elasticity. Similar predictive looking to future ball bounce locations were also reported when participants watched recorded tennis matches (Henderson, 2017).

These studies provide strong evidence that people make predictive eye movements while viewing naturalistic activity. However, no previous studies have tested whether predictive looking varies as a function of event structure. Therefore, in a series of two studies that were designed as close replications of one another, we used a new anticipatory looking task, called the Predictive Looking at Action Task (PLAT) to investigate the time course of predictability. For this task, participants’ eyes were tracked as they watched movies of an actor performing an everyday activity that consisted of sequences of goal-directed actions. Participants were not told to engage in any explicit task other than paying attention to the movie. Prediction was measured based on the amount of time participants spent looking at the object the actor was about to touch during the three seconds before the actor actually contacted the object. This task therefore allowed predictive looking to be time locked to object contact and made it possible to analyze the time course of predictive looking. We hypothesized that predictive looking to the to-be-contacted object would increase as time to object contact approached.

After watching each of the movies once, participants segmented the movies into meaningful units of activity twice: once to identify the largest meaningful units of activity (coarse events) within each movie and once to identify the smallest meaningful units of activity (fine events) within each movie. The locations at which participants identified event boundaries were time-locked to their predictive looking behavior during the passive watching condition. We hypothesized that participants would spend less time looking at the target object when object contact occurred around an event boundary compared to when object contact occurred in the middle of an ongoing event. Although this could manifest as a simple main effect, whereby predictive looking would decrease around event boundaries throughout the full three seconds before contact, predictability could also vary over time. Specifically, a decrease in predictive looking could manifest in an interaction, such that predictive looking might decrease up until it becomes clear what the actor is about to do, and then return to baseline once the actor’s actions make it obvious what the actor is about to contact.

## Materials and Methods

The present data come from a larger study investigating oculomotor control in naturalistic event viewing. A previous report (Eisenberg & Zacks, 2016) characterized the effect of event boundaries on the size and frequency of eye movements, and on pupil diameter. All of the analyses reported here are new.

### Participants

Participants for the first study were recruited from the Washington University subject pool and were either give course credit or were paid $10 for their time. Thirty-two people participated in the first study, but four were dropped from all analyses because of inability to calibrate the eye tracker (2), self-reported lazy eye (1), and self-withdrawal from the study (1). Therefore, data from 28 participants were included in the analyses reported here (50% female, age range: 18-25, mean age: 20.6). Participants for the second study were recruited from the Volunteer for Health participant registry, which is a subject pool maintained by the Washington University School of Medicine. Thirty-two participants finished the second study, but seven were dropped from all analyses because of inability to calibrate the eye tracker (5), failure to follow instructions (1), and technical error leading to loss of data. Therefore, data from 25 participants were included in the analyses for the study (68% female, age range = 22-50, mean age = 34). The Washington University Human Research Protection Office approved both studies.

### Materials

Three movies of actors performing everyday activities were used in each study. The three movies used in the first study were an actor making copies and putting together a binder (349 s), an actor sweeping the floor (329 s), and an actor changing a car tire (342 s). The movies for the second study were an actor making breakfast (329 s), an actor preparing for a party (376 s), and an actor planting plants (354 s). All six of these movies were filmed from a fixed, head-height perspective, with no pan or zoom.

### Self-Report Measure

Before beginning the eye-tracking tasks, participants completed a demographics questionnaire that included age, gender, handedness, ethnicity, foreign language knowledge, occupational history, educational history, marital status, health status, and level of typical physical activity.

### Behavioral and Oculometric Measures

Participants in both studies first watched three movies without any explicit task other than to pay close attention to the movies. After watching all three movies, participants watched the three movies two more times. During these latter two viewings, they were asked to press a button whenever they believed that one meaningful unit of activity had ended and another had begun. On one viewing, they were asked to identify the smallest units that were natural and meaningful to them (fine grain segmentation); during the other repetition, they were instructed to identify the largest units that were natural and meaningful (coarse gain segmentation). For example, a typical participant might have identified a coarse unit that could be described as “making toast,” and fine units within that coarse unit that could be described as “opening a bag of bread,” “putting bread in the toaster,” and “turning on the toaster.” (Participants were not given any such descriptions or specific instructions as to what should constitute a fine or coarse unit, beyond those given above.) The order of fine and coarse segmentation was counterbalanced across participants. In addition, participants were not told anything about the event segmentation task until after they had finished watching all three movies passively in order to ensure that participants would not covertly button press in the passive condition.

Throughout all of these tasks, gaze location from the participants’ right eye was tracked using an infrared eye tracker (EyeLink 1000; SR Research Ltd., Mississauga, ON, Canada) that sampled at 1000Hz. The eye tracker camera was mounted on the SR Research Desktop Mount. Participants were instructed to keep their heads in an SR Research chin/forehead reset throughout all of the tasks to minimize head movement during the tasks. The camera was positioned 52 cm from the top of the forehead rest. The movies were presented on a 19 in (74 cm) monitor (1400×900 resolution, viewing distance of 58 cm from the forehead rest, viewing angle of 38.6°). Data was exported from Data Viewer software (SR Research Ltd., Mississauga, ON, Canada) into text files, which were then imported into R (R Core Team, 2014) for analysis.

### Data Analysis

#### Time Course of Anticipatory Looking

The time course of anticipatory looking on the PLAT was investigated by determining how much time participants spent looking at the objects the actor was about to touch during the 3000 ms before contact. First, for each movie, an experimenter identified all of the time points at which the actor came into contact with an object. Dynamic interest areas capturing the 3000 ms before contact through 1000 ms after contact were then placed around each contacted object. Interest areas were placed using the following rules: (1) All interest areas were rectangular in shape, (2) No interest areas were allowed to overlap in time and space, (3) If potential interest areas overlapped, only the first interest area was kept, (4) If the actor contacted an object by touching it with another object, the object in direct contact with the actor was considered the object of interest (e.g., if the actor put a bowl on the counter, the bowl was considered the object of interest), (5) Only objects that were fully onscreen when contacted were considered objects of interest, (6) If the longest dimension of an object was smaller than 105 pixels (visual angle of 2.9°), the interest area was created around the entire object, and if the longest dimension of an object was larger than 105 pixels, the interest area was created around the part of the object that the actor contacted, and (7) For objects smaller than 48 pixels (visual angle of 1.3°) on any side, interest areas were created with a minimum size of 48 pixels per side. (See Fig. 1 for an example movie frame with an interest area highlighted). For the movies in the first study, there were 51, 45, and 29 dynamic interest areas for the binder, sweeping, and tire movies, respectively. For the movies in the second study, there were 49, 48, and 34 dynamic interest areas for the breakfast, party, and plants movie, respectively.

**Fig. 1.**
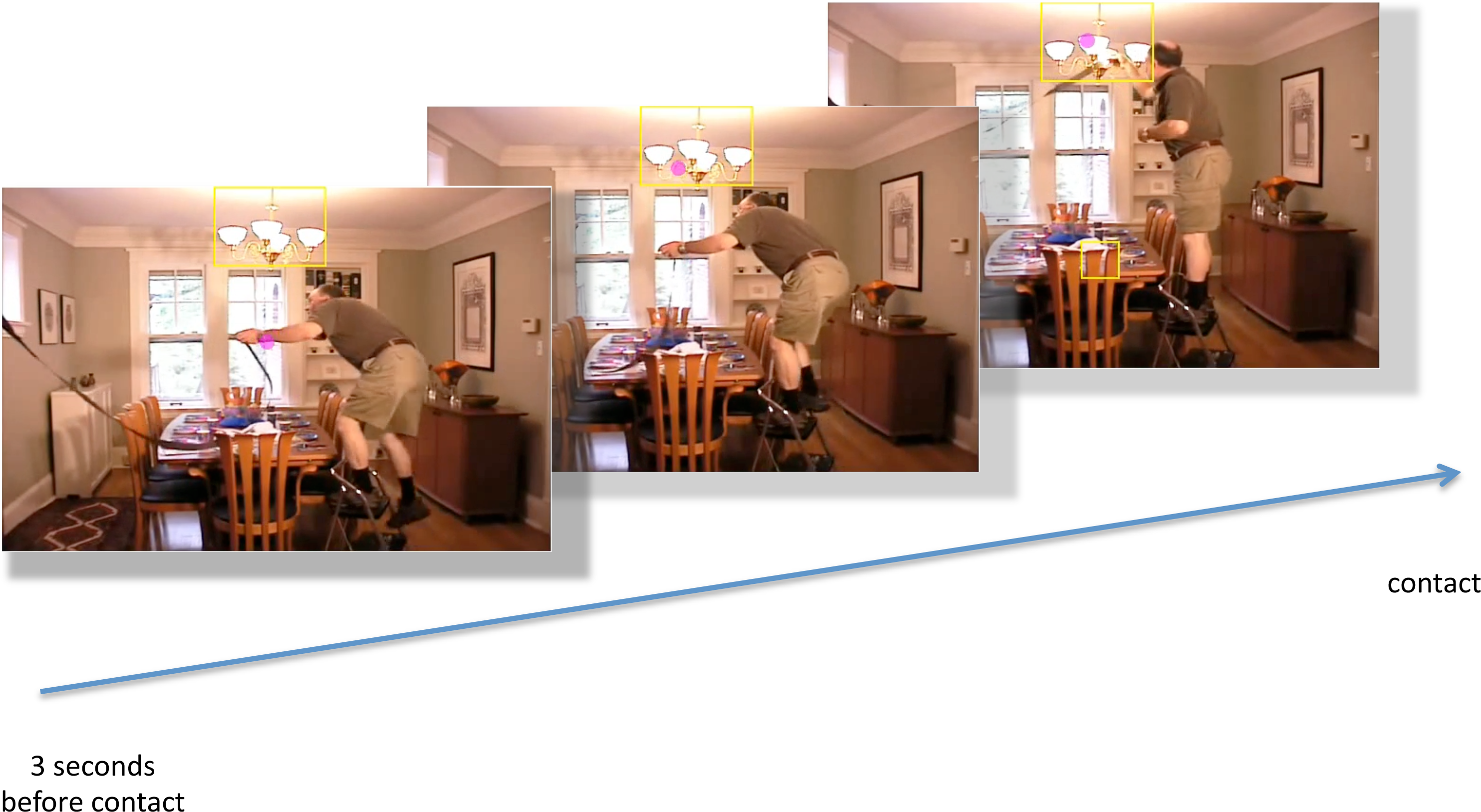
Three example frames taken from one of the movies used in study 2. The first frame is taken from around three seconds before the actor contacted the chandelier and the third frame is taken from around the time the actor contacted the chandelier. The yellow box represents the interest area, which was drawn around the chandelier—the object the actor was about to contact in order to hang a streamer. The purple dot represents the gaze location of one participant. Here, the participant looked at the chandelier before the actor contacted it. (Participants did not see the yellow box or their own gaze location while performing the task.)

Once the dynamic interest areas were identified for each movie, the eye tracking data from the three seconds before contact were divided into six 500 ms bins. Then, we calculated for each subject how long their gaze fell within the interest area for each of the six time bins. This variable was the dependent measure in the reported mixed-effects model analyses. To determine how predictability varied as time to object contact approached, the lme4 and lmerTest packages in R (Bates, Maechler, Bolker, & Walker, 2015; Kuznetsova, Brockhoff, & Christensen, 2014) were used to compare nested mixed-effects models to determine whether including the fixed effect of time bin explained significant additional variance in the dependent variable. The *effects* package in R (Fox, 2003) was used to estimate the fixed effects for plotting and to calculate confidence intervals for the fixed effects in the linear mixed effects models.

#### Time Course of Predictive Looking around Event Boundaries

To investigate the time course of predictability around event boundaries, we coded whether each object contact happened within 1000 ms of a coarse boundary or within 1000 ms of a fine boundary. This was done separately for each participant, based on the segmentation data from their subsequent viewing of the movie. (Only the initial viewing eye tracking data were analyzed, to exclude contamination from the cognitive operations necessary for the segmentation task.) Mixed-effects models were analyzed using the lme4 and lmerTest packages in R (Bates, Maechler, Bolker, & Walker, 2015; Kuznetsova, Brockhoff, & Christensen, 2014) to determine whether there was a main effect of being near a fine or coarse event boundary on predictive looking, and whether this interacted with time relative to contact. (If the temporal window before object contact were sufficiently large, one would expect no effect for the earliest timepoints, before predictive looking is possible, with the effect building as time to contact approached.) Again, the *effects* package in R (Fox, 2003) was used to estimate fixed effects and confidence intervals. For the first study, 4.1% percent of the interest areas were classified as being near a coarse event boundary, 41.1% were classified as being near a fine event boundary, and 15.2% were classified as being near both a coarse and a fine boundary. For the second study, 5.5% were classified as being near a coarse event boundary, 34.8% were classified as being near a fine event boundary, and 17.6% were classified as being near both a coarse and a fine boundary.

## Results

### Time Course of Predictability

To investigate how predictability varied over the three seconds before contact, nested mixed-effects models with item, movie, and subject included as random effects and with and without time bin included as a fixed effect were tested. For both studies, including the fixed effect of time bin in the model explained significant additional variance in the amount of time participants spent looking at the target object (Study 1: χ^2^ = 2183.1, df = 5, p < .001; Study 2: χ^2^ = 1858, df = 5, p < .001), suggesting that predictability increased as object contact approached. Figure 2 displays the amount of time participants spend looking at the target object within each of the six 500 ms bins during the three seconds before the actor contacted the target object.

**Fig 2:**
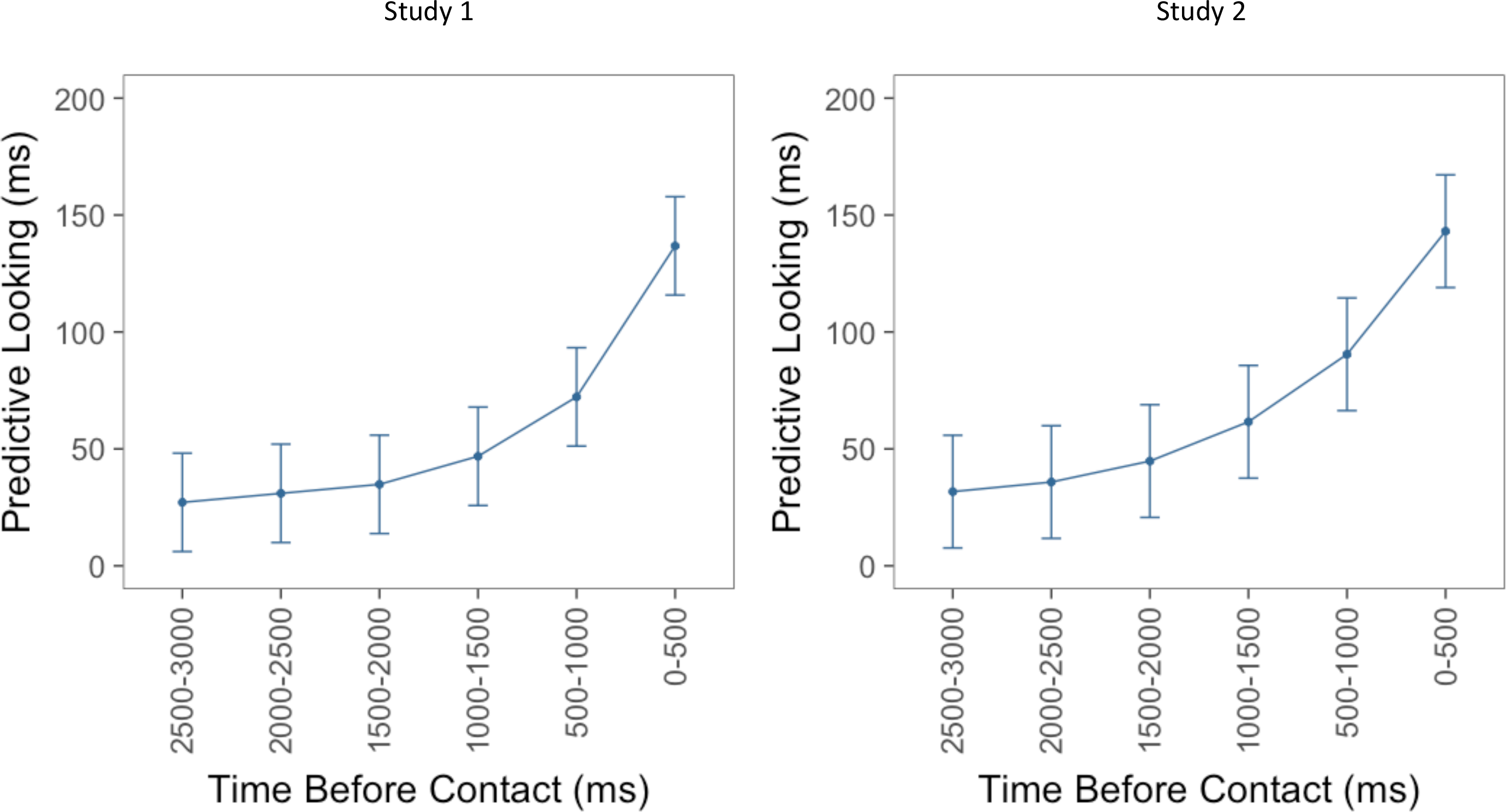
The panel on the left displays the results of the linear mixed effects model for the first study. The y-axis displays the beta values representing the amount of time participants spent looking at the target object during each of the six 500 ms bins. The panel on the right displays the same information for the second study. The x-axis for both figures displays the six bins, where the leftmost bin on each x-axis represents the time farthest away from contact and the rightmost bin represents the time closest to contact. Error bars depict 95% confidence intervals.

### Time Course of Predictability Around Event Boundaries

To investigate the time course of predictability around event boundaries, mixed-effects models were tested with time bin and boundary type (no boundary, fine boundary, coarse boundary, both fine and coarse boundaries) as fixed effects and item, movie, and subject as random effects. For both studies, there was a significant main effect of bin (Study 1: F = 233.72, df = 5, p <. 001; Study 2: F = 211.75, df = 5, p < .001), and a significant interaction between time bin and boundary type (Study 1: F = 2.26, df = 15, p = .004; Study 2: F = 1.69, df = 15, p = .04). The form of the interaction is illustrated in Figure 3 and Figure 4: For objects contacted in the middles of events participants looked to the object relatively early, whereas for objects contacted near event boundaries they tended to look more just before object contact. The main effect of boundary type was not significant (Study 1: F = .95, df = 3, p = .42; Study 2: F = 1.38, df = 3, p = .25). To determine whether boundary types differed significantly from one another, three nested models were tested: a null model containing a binary variable coding whether there was an event boundary present or not, a model with this binary variable and a variable coding for the effect of fine boundaries, and a model adding a variable coding for the effect of coarse boundaries. All three models also included interaction terms coding for the interaction of time point and boundary type. None of these models were significantly different from one another (Study 1: largest χ^2^ = 12.86, df = 12, p = .38; Study 2: largest χ^2^ = 13.41, df = 12, p = .34). Therefore, all three boundary conditions (fine, coarse, and both fine and coarse) were collapsed into a single boundary variable, as depicted in Figure 4. There was a significant main effect of time bin (Study 1: F = 422.05, df = 5, p < .001; Study 2: F = 370.82, df = 5, p < .001), and again there was a significant interaction between time bin and boundary type for both studies (Study 1: F = 4.75, df = 5, p < .001; Study 2: F = 2.92, df = 5, p = .01). The main effect of boundary was again not significant (Study 1: F = 0.13, df = 1, p = .72; Study 2: F = 1.50, df = 1, p = .22). To assess which, if any, individual time points had significant differences between the boundary and within-event conditions, we fitted mixed-effects models testing the difference for each time point. None of these were significant (Study 1: largest F = 3.03, p = .08; Study 2: largest F = 1.04, p = .31).

**Fig. 3:**
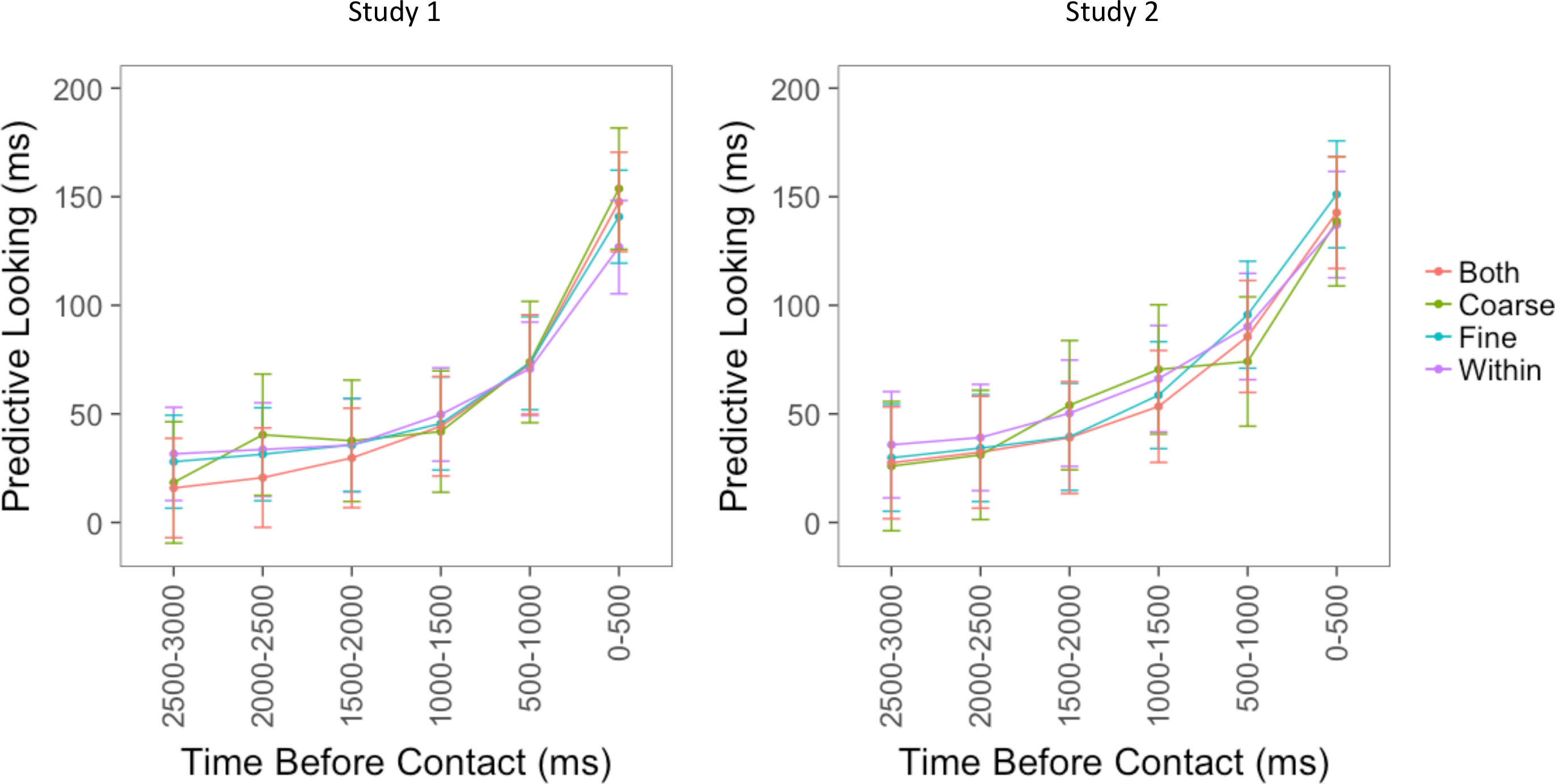
The panel on the left displays the results of the linear mixed effects model for the first study. The y-axis displays the beta values representing the amount of time participants spent looking at the target object during each of the six 500 ms bins. The panel on the right displays the same information for the second study. The x-axis for both figures displays the six bins, where the leftmost bin on each x-axis represents the time farthest away from contact and the rightmost bin represents the time closest to contact. Each colored line represents the amount of time participants spent looking at the target object for each boundary type. Error bars depict 95% confidence intervals.

**Fig. 4:**
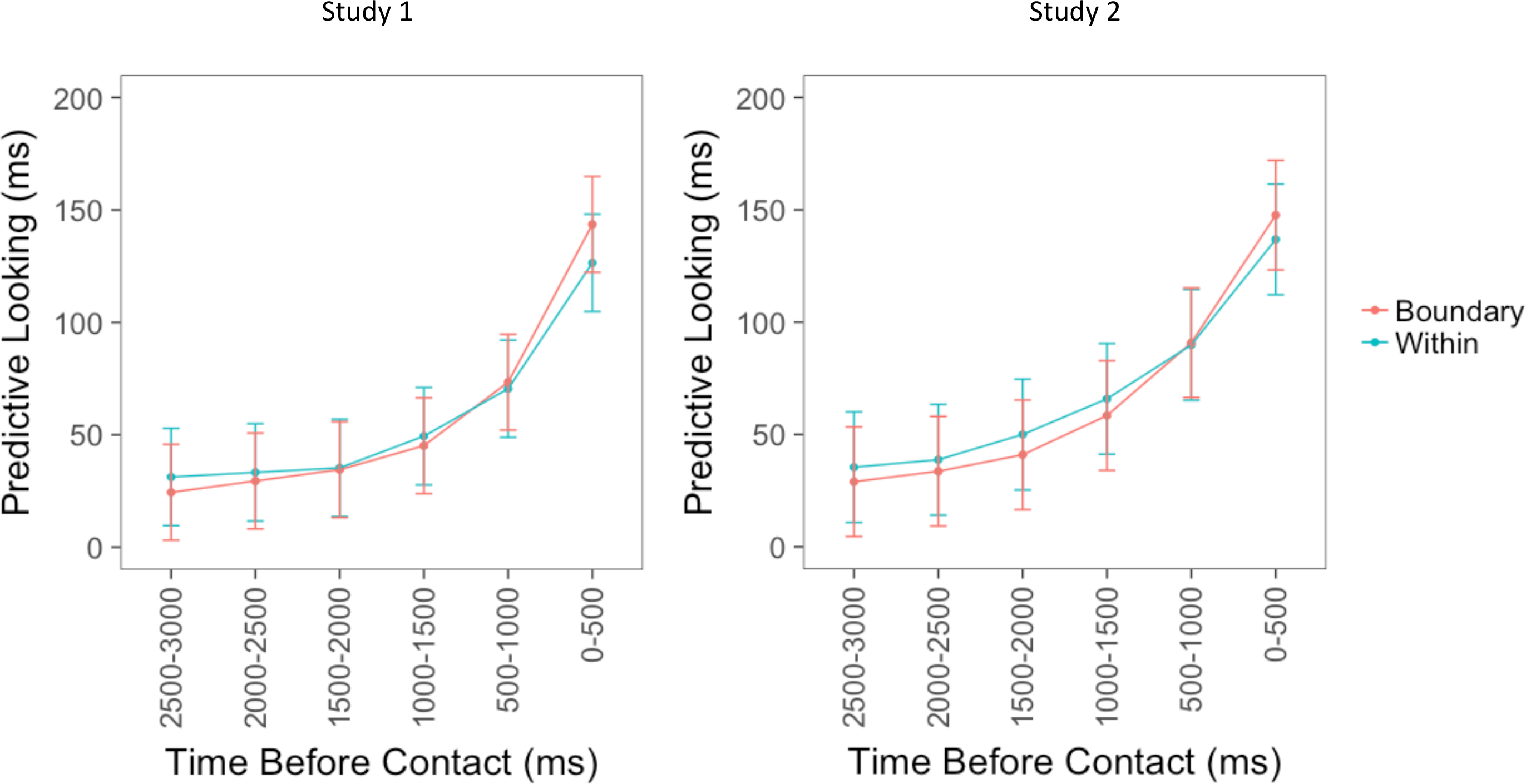
The panel on the left displays the results of the linear mixed effects model for the first study. The y-axis displays the beta values representing the amount of time participants spent looking at the target object during each of the six 500 ms bins. The panel on the right displays the same information for the second study. The x-axis for both figures displays the six bins, where the leftmost bin on each x-axis represents the time farthest away from contact and the rightmost bin represents the time closest to contact. The red line displays predictive looking collapsed across boundary types (fine, coarse, both fine and coarse). The blue line displays predictive looking when there were no event boundaries nearby (within events). Error bars depict 95% confidence intervals.

In sum, both experiments showed an interaction between time and boundary condition, such that around event boundaries compared to within events, participants engaged in less predictive looking earlier on and engaged in similar or more predictive looking during the 500 ms before object contact. There was no evidence that coarse and fine boundaries differed from each other, and in neither experiment could the effect be statistically localized to any individual time points.

## Discussion

The current studies introduce the PLAT as a tool for investigating the time course of predictive looking and provide the first demonstration of the dynamics of predictive looking during viewing of a naturalistic sequence of activities. In both studies, the amount of time participants spent looking at a target object increased as the actor came closer to contacting the object. This result provides validation that this looking behavior can be used as a noninvasive measure of prediction during ongoing comprehension.

After determining that the PLAT could be used to investigate predictive processing, it was used to study the dynamics of predictability around event boundaries. In both studies, the amount of time participants spent looking at the object the actor was about to contact increased progressively during the three seconds before contact. In addition, there was a significant interaction such that participants engaged in less predictive looking around event boundaries than within events for time bins farther away from contact and engaged in more predictive looking around event boundaries than within events for time bins closer to contact. All of the results replicated across the two studies, providing strong evidence for these effects.

Our interpretation of this interaction is that in the middle of an event it is easier to predict what will happen two to three seconds ahead, so viewers’ eyes are somewhat likely to jump ahead to the to-be-contacted object. For example, in Figure 1, once the actor steps on the ladder and reaches up toward the chandelier with the streamer in his hand, it is fairly clear which object the actor is about to touch because there are no other objects nearby. When this happens, their eye may have moved on by the time the actor’s hand actually reaches the object, leading to decreased looking times right before contact. In contrast, near an event boundary it is more difficult to predict two to three seconds ahead, so the eye is more likely to reach the object being manipulated just before the actor’s hand arrives, leading to increased looking times right before contact. These effects are not huge but they appear robust.

The two studies described here were purposely designed as close replications of one another in order to determine the reproducibility of the results. However, while the design of the studies was nearly identical, the populations and movies used in the studies differed. In the first study, participants were undergraduates recruited from the university’s participant pool, whereas in the second study, the participants were recruited from the general population of St. Louis and included a much more diverse age range. In addition, completely different movies were used in the two studies to ensure that the results were not specific to particular sequences of naturalistic activity, but instead were generalizable to other sequences of actions. The pattern of results from the two studies was almost identical, providing strong evidence for the findings reported here.

These results, and this interpretation, are consistent with previous studies investigating prediction around event boundaries using explicit measures. In three studies, Zacks, Kurby, Eisenberg, and Haroutunian (2011) asked participants to watch similar movies of everyday activities to those used in the present studies. Each movie was paused eight times, four times around event boundaries and four times within events and participants either made forced choice two-alternative decisions about what would happen five seconds later in the movie or they made yes-no decisions about whether one image would appear five seconds later in the movie. The authors found that participants were more accurate in making predictions when the movies were paused within events than when they were paused right before event boundaries.

We initially predicted that the presence of an event boundary would be associated with lower predictive looking overall, a main effect, in addition to the interaction observed. One possibility is that this main effect would have been observed if we had looked farther back in time before each object. As can be seen in Figures 3 and 4, the largest difference between the conditions appears to be at the earliest timepoints. In the explicit prediction study of Zacks et al. (2011) a difference was found for predictions of five seconds in the future. However, a forced choice task is very different than making open-ended predictions by looking around a visual space. Therefore, it is possible that prediction would be equally bad around event boundaries and within events as object contact becomes farther away in time, especially since participants in the current study spent an average of less than 50 milliseconds looking at the target object from 2500 to 3000 milliseconds before contact.

It is also possible that the lack of a main effect of boundary type is due in part to the large difference in the number of observations for each boundary condition. As noted above, only 4.1 and 5.5 percent of the observations occurred near coarse boundaries, which likely explains the large confidence intervals for this condition. However, this difference in number of observations cannot fully explain the results. First, the results presented above were obtained using linear mixed effects modeling, which took into account the different numbers of observations among the conditions. In addition, when the boundary conditions were collapsed into within events versus around event boundaries, the number of observations in each condition were more similar (60.4 and 57.9 percent of observations were around event boundaries for the two studies).

Therefore, it is unlikely that additional observations for the coarse boundary condition would have dramatically altered the results.

In addition to its utility in the current studies, the PLAT has strong potential for utilization in other studies investigating predictive processing. The PLAT allows for the collection of large amounts of data in a short amount of time, as each of the five to six minute movies contained between 29 and 51 target objects and three thousand data points were analyzed for each of these trials. Although this study was not designed to investigate individual differences in predictive looking, the large amount of data that can be collected using the PLAT positions it as a potentially powerful individual differences measure. If individuals vary in their time course of predictive looking, performance on the PLAT might correlate with other cognitive measures such as working memory or executive function. It would be informative to determine whether predictive looking behavior is fully explained by other cognitive abilities or whether it is a separate cognitive ability in a similar way as working memory is at least partially independent of executive processing.

In addition to its utility in studying prediction under naturalistic comprehension situations, the PLAT has potential for studying predictive processing in populations that are unable to perform explicit prediction tasks. For example, the task can be used with infants or very young children, who would not be capable of performing an overt prediction task. The task could also be used to investigate predictive processing in clinical populations who may not have the verbal or motor ability to complete a prediction task that requires overt responses.

## Conclusion

People engage in prediction in almost every moment of the day, and understanding how predictability varies over time is integral to understanding how people comprehend ongoing activity. Using a novel predictive looking task, the two current studies extended previous research on the time course of predictability, finding that participants engaged in less predictive looking around event boundaries than within events at time points furthest from contact and participants engaged in more predictive looking around event boundaries immediately before contact. These results are consistent with previous studies that found decreases in predictability at times of greatest change in the environment. The two current studies also demonstrated the utility of the PLAT as a sensitive measure of the time course of prediction, and the PLAT can easily be extended to naturalistically study prediction ability in healthy adults, clinical populations, and even infants and young children.

## List of Abbreviations

EST: Event Segmentation Theory
PLAT: Predictive Looking at Action Task

## Ethics Approval and Consent to Participate

All participants provided informed consent to participate in the study, and the Washington University Human Research Protection Office approved both studies.

## Consent for Publication

Not Applicable

## Availability of Data and Materials

All data and materials are freely available from the authors of the manuscript on written request.

## Author Contributions

Michelle Eisenberg had the primary role in study design, overseeing data collection, conducting analyses of the data, interpreting the data, and writing this manuscript. Jeffrey Zacks also had an integral role in study design, interpreting the data, and writing this manuscript. Shaney Flores developed the guidelines for identifying interest areas and applied these guidelines to identify the interest areas for the movies included in this study. He also played an integral role in the data collection process.

## Acknowledgements

The authors thank Mobolaji Fowose, and Angela Lee for their help running participants.

